# Neuromagnetic activation dynamics of stimulus-locked processing during a naturalistic viewing

**DOI:** 10.1101/711457

**Authors:** Adonay S. Nunes, Nataliia Kozhemiako, Alexander Moiseev, Robert A. Seymour, Teresa P. L. Cheung, Urs Ribary, Sam M. Doesburg

## Abstract

Naturalistic stimuli such as watching a movie while in the scanner provide an ecologically valid paradigm that has the potential of extracting valuable information on how the brain processes complex stimuli in a short period of time. Naturalistic viewing is also easier to conduct with challenging participant groups including patients and children. Given the high temporal resolution of MEG, in the present study, we demonstrate how a short movie clip can be used to map distinguishable activation dynamics underlying the processing of specific classes of visual stimuli such as face and hand manipulations, as well as auditory stimuli with words and non-words.

MEG data were collected from 22 healthy volunteers (6 females, 3 left handed, mean age – 27.7 ± 5.28 years) during the presentation of naturalistic audiovisual stimuli. The MEG data were split into trials with the onset of the stimuli belonging to classes of interest (words, non-words, faces, hand manipulations). Based on the components of the averaged sensor ERFs time-locked to the visual and auditory stimulus onset, four and three time-windows, respectively, were defined to explore brain activation dynamics. Pseudo-Z, defined as the ratio of the source-projected time-locked power to the projected noise power for each vertex, was computed and used as a proxy of time-locked brain activation. Statistical testing using the mean-centered Partial Least Squares analysis indicated periods where a given visual or auditory stimuli had higher activation. Based on peak pseudo-Z differences between the visual conditions, time-frequency resolved analyses were carried to assess beta band desynchronization in motor-related areas, and inter-trial phase synchronization between face processing areas. Our results provide the first evidence that activation dynamics in canonical brain regions associated with the processing of particular classes of visual and auditory stimuli (words, faces, etc.) can be reliably mapped using MEG during presentation of naturalistic stimuli. Given the strength of MEG for brain mapping in temporal and frequency domains, the use of naturalistic stimuli may open new techniques in analyzing brain dynamics during ecologically valid sensation and perception.

**Highlights:**

- A time-locking analysis was employed in naturalistic stimuli paradigm.
- Specific visual and auditory stimuli from the movie were mapped in brain space.
- Motor β-suppression was evident in periods of watching hand manipulation.
- Increased synchronization between core face-processing areas was found around 200 and 300ms in the face condition.
- Naturalistic viewing paradigms provide a reliable approach for investigating brain dynamics.

## 1. INTRODUCTION

Traditional neuroimaging paradigms typically sacrifice ecological validity for high experimental control. Such approaches ensure that observed brain effects can be attributed to specific experimental manipulations with high confidence. However, brain processes studied under this type of paradigm might lack transferability to real life conditions (Schmuckler, 2001). For example, real life sensory processing and behavior is complex and dynamic involving interactions of many context-dependent stimuli through different sensory modalities which are constantly changing, whereas traditional neuroimaging studies often isolate a specific task with highly constrained and simplified stimuli which are repeated several hundreds of times to obtain reliable brain activity patterns. An alternative approach is a naturalistic paradigm which consists of more complex and dynamic stimuli using speech, videos and/or music. Such paradigm also help to increase the compliance of subjects (Centeno et al., 2016; Vanderwal et al., 2015) and allow the recording of brain activity from populations that may not be able to perform repetitive tasks involving hundreds of trials and behavioral responses, such as children or individuals with disorders (Vanderwal et al., 2019).

Since Hasson et al., 2004 demonstrated that brain activity is synchronized across subjects while watching movies, intersubject correlation (ISC) analysis of fMRI data has become increasingly used and currently available in major fMRI datasets (HCP (Glasser et al., 2013) CamCan (Taylor et al., 2017). ISC has provided evidence of brain activity at different frequencies involved in naturalistic stimuli processing (Kauppi et al., 2010), shared brain activations during story comprehension across different languages (Honey et al., 2012), mapped the similarities in brain responses in people with similar interpretation of the naturalistic stimuli (Nguyen et al., 2019), and identified brain areas related to successful episodic memory formation (Hasson et al., 2008) and emotion processing (Nummenmaa et al., 2014, 2012), as well as characterizing movie-induced functional connectivity changes (Demirtaş et al., 2019; Simony et al., 2016).

The approach of ISC has been also applied to MEG data and revealed synchronization across subject consistent with fMRI findings (Lankinen et al., 2018). The high temporal resolution of MEG allowed extending the framework of ISC to different frequencies and it has been shown more pronounced ISC at low (<10 Hz) frequency bands (Lankinen et al., 2014). However, previous studies focused on approaches that relied on adapting the MEG time series, such as multiset canonical correlation analysis, or smoothing the brain activity across vast anatomical areas, in order to reduce the intrinsic noise of MEG signals (Chang et al., 2015; Lankinen et al., 2014). This, however, seems to limit the ability to exploit the richness of MEG signal during sensory and perceptual processing of naturalistic stimuli.

In this study, we use a classical approach such as time locking analysis, but employed within the context of a novel naturalistic stimuli paradigm, to test the hypothesis that stimulus-locked MEG activity presented in naturalistic stimuli can reliably map and differentiate brain dynamics associated with particular classes of visual and auditory stimuli. Specifically, we mapped and contrasted naturalistic faces with hand manipulation stimuli, as well as contrasting word and non-word auditory stimuli. We present a novel framework that involves time locking signal to the onset of movie visual and auditory events. We use the ratio of beamformer projected time locked power by projected noise (Pseudo-Z) to localize time locked activity related to face and hand movements stimuli and to words and non-words sounds. Based on spatial mapping in the visual conditions, we further tested the effect of β-suppression induced while observing hand movements, and phase locking between the areas that were more activated while watching movie contents with faces. To our knowledge, this is the first time such dynamical approach with milliseconds time resolution is applied to this type of paradigms, allowing to find areas with higher signal-to-noise ratio (SNR) associated with specific movie content.

## 2. MATERIALS & METHODS

### 2.1. Participants

MEG data were collected from 22 healthy participants (6 females, 3 left handed, mean age – 27.7 ± 5.28 years). All participants had normal or corrected to normal vision. None of the participants reported hearing problems or any medical condition including neurological disorders. Informed consents were obtained from all participants. The present study was approved by the Research Ethics Board of Simon Fraser University and the Fraser Heals Research Ethics Board.

### 2.2. MEG and MRI data collection

MEG data were collected at a sampling frequency of 1200 Hz in a magnetically shielded room using a 275 channel MEG system (CTF systems; Coquitlam, Canada). The data recording was performed while the participants where in supine position viewing a short movie clip on a screen 40 cm above their eyes. The clip was presented two times for each participant with a short break in between. The audio from the movie clip was presented through MEG-compatible earphones and prior to the data collection a short test was conducted to ensure that a participant could hear the sound from both earphones. Participants were instructed to stay as still as possible and pay attention to both movie presentations. Head position in the dewar was continuously tracked using three fiducial coils attached at the nasion and left and right preauricular locations. Prior to the MEG data collection, the head shape of each participant was digitalized using Polhemus FASTRAK digitizer for co-registration of MEG data with the anatomical MRI. The T1-weighted structural images (Philips 3T Ingenia CX) were collected for each participant (TE = 3.7ms, flip angle = 8°, FOV = 256 × 242, matrix size = 256 × 242, slice thickness = 1 mm, number of slices = 213, sagittal orientation).

### 2.3. Natural viewing paradigm

Presented movie clip was a compilation of segments from the movie “*Charlie and the chocolate factory”, 2005* directed by Tim Burton with a total duration of 4 minutes and 32 seconds (Fig.1A). The changes between scenes were marked by a photodiode dot on the first frame of each new scene. For the analysis, each movie frame was manually classified into the following categories: faces (56 occurrences), hand manipulations (42 occurrences), distant bodies (28 occurrences), non-human objects and scenery views (9 occurrences) [Fig. 1B]. The sound wave from the movie was also manually classified using audioLabeler (MATLAB 2019a) into three categories: word sounds with clear onset (95 occurrences), non-word sounds with clear onset (59 occurrences), background noise (88 occurrences). The timings of stimuli occurrences combined from both movie presentations were used for data selection to investigate sensory processing as described in the data analysis section.

**Fig.1.**
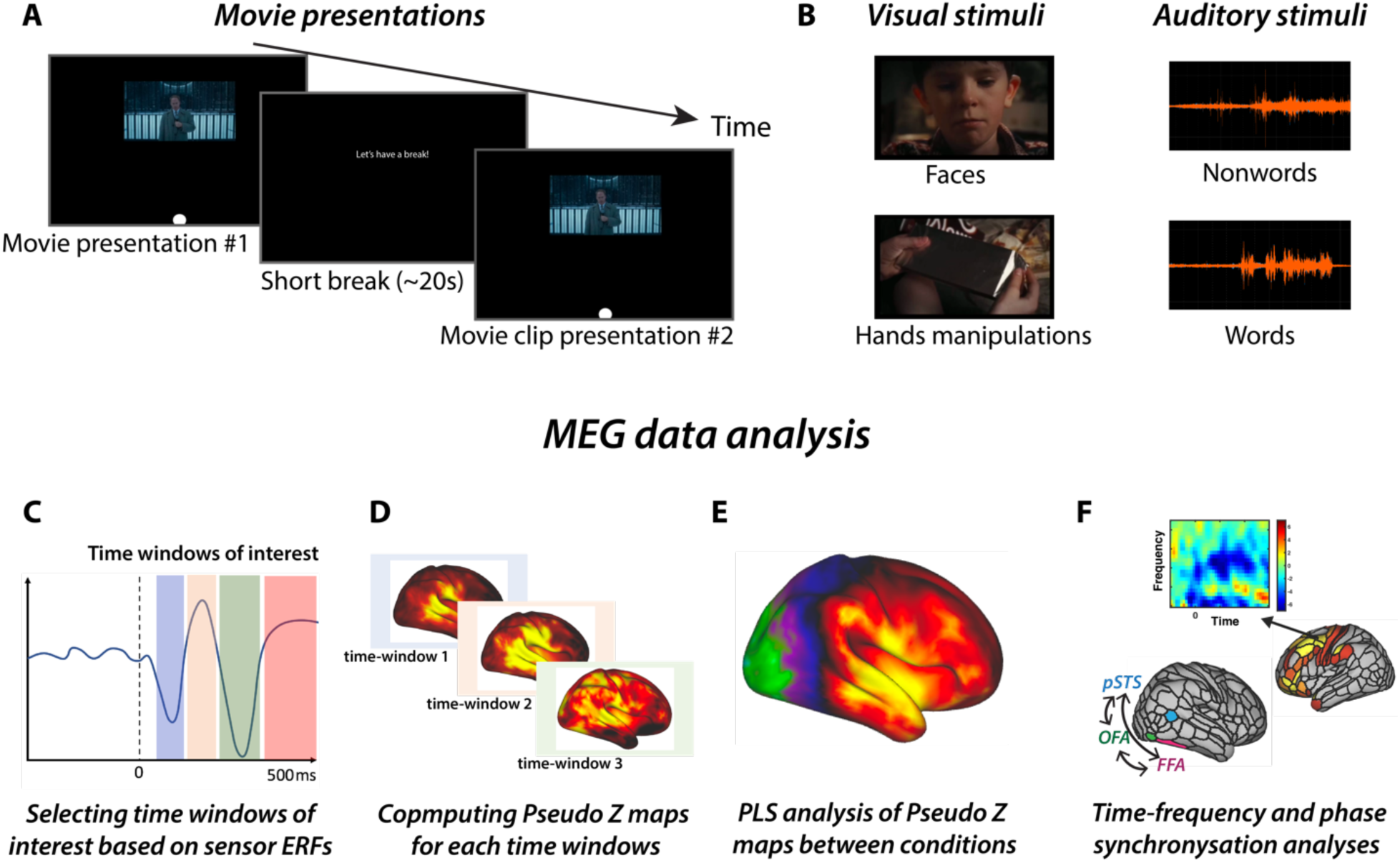
Naturalistic stimuli presentation and analysis workflow: A – Presentation of a movie clip (4 min 32 sec) twice with a short break of about 20 sec in between; B – Manual classification of visual and auditory stimuli into subcategories; C – Sensor level event-related fields used to define time windows of interest for source analysis; D – Computation of Pseudo-Z maps for selected time windows; E – Statistical analysis of Pseudo-Z maps differences between different conditions; F – time-frequency and connectivity analysis between different conditions for specific brain areas.

### 2.4. Data analysis

First, the segments of data with head movement exceeding 5 mm in any directions for any of the three fiducial coils were detected and excluded from the analysis. Only five participants exceeded this threshold and even for those participants, the maximum length of removed data was less than 3 sec for the whole recording. The data was band-pass filtered from 1 to 150 Hz and a 60 Hz notch-filter was applied to remove line noise. Then Independent Component Analysis was performed to define and subsequently exclude the independent components composed of EOG (blinks, saccades) and heart artifacts using Fieldtrip toolbox (Oostenveld et al., 2011). For every occurrence of a stimulus, two seconds long epoch (trial) was extracted starting 500ms before the stimulus onset.

#### 2.4.1. Sensor level event-related fields

To investigate event related responses to visual (faces and hands manipulations) and auditory (words and non-words) stimuli on the sensor level, the data was time locked to the stimulus onset and averaged across the trials using Fieldtrip toolbox (Oostenveld et al., 2011). The data were filtered from 1 to 40 Hz and baseline corrected (the baseline was defined as 500ms before the stimulus onset). Subsequently, the trials were averaged across all participants for each of the conditions, i.e. visual-faces, visual-hand movements, auditory-words, auditory-non-words. We used the event-related fields to define time windows of interest for the source-based analysis.

#### 2.4.2. Source modeling and forward solution

For each subject, we segmented their MRI to extract the cortical surface. A cortical mesh of 4K vertices on a standard space was then constructed and used as source locations. A single-shell headmodel was created for each subject based on the subject’s inner skull. The forward solution was computed for each vertex from the cortical mesh using Fieldtrip toolbox (Oostenveld et al., 2011).

#### 2.4.3. Source localization

To localize time-locked activity, a scalar beamformer (Robinson and Vrba, 1999) was used to find the Pseudo-Z, which is the ratio of the projected source power to the projected noise power for a given location on the cortical mesh, defined as:

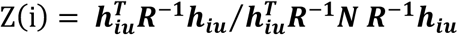

where **h** is the forward solution at location *i* with orientation **u, R** = ⟨***bb***^T^⟩ is the signal covariance of M sensor time series **b** collected over intervals of interest, and **N** is the MxM noise covariance collected over some intervals with no current stimuli present as explained below; ⟨•⟩ denotes statistical averaging and the superscript ‘T’ matrix transposition. A higher SNR can be obtained with event-related evoked activity, as is our case, where responses were time-locked to the onset of movie contents. The event-related localizer is obtained as:

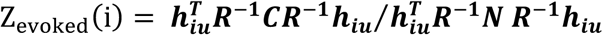

Here **C** denotes the 2^nd^ moment matrix of the trial-averaged sensor time series 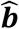, time-averaged over an interval of interest *t* time points long: 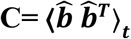.

For each subject, the spatial Pseudo-Z distribution was transformed to z-scores by subtracting the mean and normalizing on the standard deviation computed over all voxels for each time window, to decrease dependence of the group results on differences in dynamic ranges of individual datasets (Moiseev et al., 2015, 2011; Nunes et al., 2019).

Here, we use Pseudo-Z as a proxy for brain activation. It reflects the ratio of evoked source power at a given time window to the source power at the times of no interest. For each visual and auditory condition at each time window, Pseudo-Z spatial distribution was computed. The Pseudo-Z calculation for the face condition, signal covariance matrix was obtained by including all trials categorized as faces (102 occurrences), for the hand manipulations were used all the trials classified as hand manipulations (84 occurrences). The noise covariance was computed based on all other trials including distant bodies and non-human objects and scenery views (56 and 18 respectively, 74 in total). Similarly, for the words condition, we used all the words trials with a clear onset (190 occurrences) to compute the signal covariance matrix and for the non-words condition, all the non-word sounds with a clear onset (118 occurrences). To compute noise covariance for both auditory trials, we used the trials classified as background noise (176 occurrences). Then, the Pseudo-Z spatial distributions were compared between categorical conditions to find peak locations that characterize face, hand-movements, words and non-words processing.

#### 2.4.4. Source reconstruction

Beamformer weights **w** were derived for each time window to optimize the spatial filtering of the beamformer at the specific time-locked window. Given the narrow time windows, to obtain robust estimates of the inverse covariance, broad band sensor time series were used. Dipole orientation **u** was selected as the optimal orientation that maximizes the evoked projected power to the projected noise by solving the generalized eigenvalue problem, and used to compute

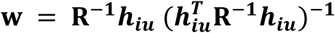

Due to computational limitations, we reduced data dimensionality by estimating an areal ***w***_***a***_ using the areas from the Multimodal Parcellation Atlas (Glasser et al., 2016), (Schoffelen et al., 2017; Seymour et al., 2018). First, spatial filters ***W***_***i***_ at locations *i* were estimated for each vertex within an area. Then singular value decomposition (SVD) was used to decompose the product of 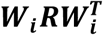. The first left singular vector ***uv***_1_, expressing most of the power covariance of the vertex within a parcellation area, was used as ***w***_***a***_ = ***uv***_**1**_***W***_***i***_. Given that the source power is center bias, the time series were normalized by dividing them by 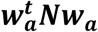. To correct for sign shifting across participants, we computed the dot product of the subject’s time series for a given area and the ***uv***_**1**_of all the subject’s areal time series. These source estimates where then used to compute ERF, connectivity and time-frequency analysis.

#### 2.4.5. Source space time-frequency (TF) analysis

Previous studies have demonstrated event related desynchronization (ERD) in alpha (8-12Hz) and beta (12-25hz) bands during hand movements or observation of hand movements performed by others (Erbil and Ungan, 2007; Puzzo et al., 2011). To further explore the perceptual dynamics caused by hand movements observation while watching a movie, we performed a TF analysis comparing faces and hand-movements. The areal time series from the areas classified in Glasser et al. (2016) as somatosensory, supplementary motor and premotor areas were selected (20 areas). Using a sliding Hanning window of 400ms, the signals were spectrally decomposed and power was estimated at 50ms steps from 6 to 30 Hz between −0.5 to 1.5 seconds from the stimuli onset. Then, the TF were baseline corrected by subtracting the mean baseline values across frequencies. The TFs for all the motor related areas between faces and hand movements were tested together for reliable statistical significance using PLS analysis.

#### 2.4.6. Source space inter-trial phase locking

Phase synchronization across time points between signals was assessed for the face and hand movements conditions between areas of the so called ‘core’ system involved in face processing (Haxby and Gobbini, 2011), namely, the occipital face area (OFA), posterior superior temporal sulcus (pSTS) and fusiform face area (FFA). To do so, spectral decomposition of the three areal time series over time was performed using a continuous wavelet transform centered at 15 frequency bins logarithmically spaced from 4 to 45 Hz from −0.1 to 0.6 seconds from the stimuli onset. Sample Phase Locking Value (sPLV) between signal pairs was computed as:

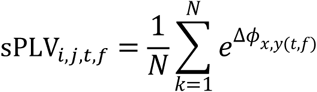

where *N* is the number of trials, Δ*ϕ* is the phase difference between *x* and *y* at time *t* and *f* frequency. As in the TF analysis, sPLV was baseline corrected to remove non-task related spurious synchronization. Then, the sPLV between the three areas of interest were tested together for significance using PLS analysis.

### 2.5. Statistical analysis

In our study, we used mean-centered Partial Least Squares (PLS) analysis for testing statistical significance of the differences between the conditions (Lobaugh et al., 2001; McIntosh and Lobaugh, 2004). It is a multivariate statistical approach that uses singular value decomposition to extract latent variables which express the most of the variance in the data. Each latent variable is composed of (i) a vector expressing the contrast between conditions, (ii) singular value ultimately representing the amount of variance explained by the latent variable, and (iii) a vector of saliences expressing the contribution of each data feature (in this study, each vertex for Pseudo-Z analysis or the combination of an area at specific timepoint at specific frequency for TF or connectivity analysis) to the contrast between conditions. Subsequently, the series of global and local tests are performed to delineate statistical significance of the latent variable. The global test is based on a series of random permutations across subjects across different conditions. It renders a single p-value reflecting the statistical significance of the contrast between the conditions. The local tests are based on bootstrapping subjects within conditions resulting in the measures that can be interpreted as z-score for each data feature. This essentially expresses each data feature’s contribution to the overall contrast between conditions. Positive or negative z-scores of 2.5 or −2.5 were chosen as thresholds since they approximately correspond to the 99% confidence interval (McIntosh and Lobaugh, 2004). Positive z-scores represent vertices with higher activation in condition A in contrast to condition B (e.g. face > hand manipulations) and vice-versa for negative z-scores (e.g. hand manipulations > face). For visualization purposes, we split z-scores into positive and negative and flip the negative z-scores, to demonstrate higher activation in one condition compared to the other with the same 2.5 threshold.

For statistical comparison of Pseudo-Z maps between conditions, we ran separate PLS analysis for each time window (four PLS analyses between face trials and hand manipulation trials; three PLS analyses between word trials and non-word trials). Given that we ran separate PLS analysis for each time window for visual and auditory trials, p-values for each PLS analysis were Bonferroni corrected by the number of PLS analyses for each condition. We also ran two separate PLS analyses to test the significance of differences between the conditions in spectral power dynamics in motor-related areas and inter-trial phase synchronization within the areas of the ‘core’ network of face processing. For all PLS analyses, we used 5000 permutation and 5000 bootstrapping rounds.

## 3. RESULTS

### 3.1. Activation dynamics during viewing of faces and hand movements

First, we investigated event related components time-locked to the visual stimuli onset which was defined as the first frame of a scene with faces or hand manipulations appearing on the screen. In Fig. 2A, the ERF responses for these two stimulus types are presented averaged across right temporal-occipital MEG channels to target the areas reported to be involved in the processing of similar stimuli (Meeren et al., 2013). The averaged ERFs were composed of well-documented components elicited by visual stimuli. Based on these waveform components four time windows were selected for the source analysis: 50-120ms, (M100 component) 120-190ms (M170 component), 190-330ms (M250 component) and 330-450ms (M400 component) which are consistent with previous studies exploring evoked brain responses to visual stimuli in standard paradigms (Puce et al., 2013).

**Fig. 2.**
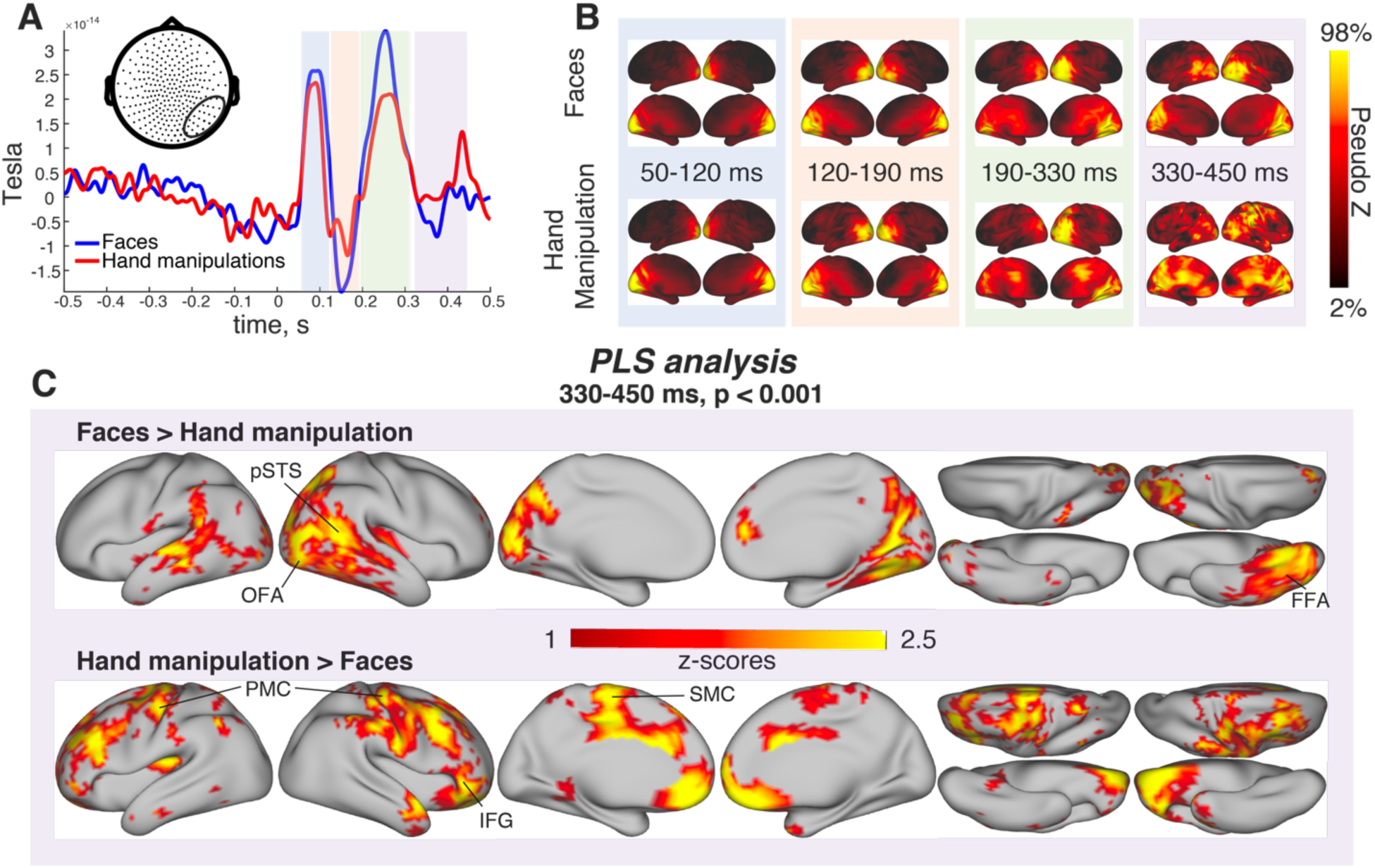
Differences in activation dynamics between different visual stimuli – frames with faces and hand manipulations presented during the movie: A – averaged across right temporal-occipital sensors event related fields for both conditions. B – Pseudo-Z maps for each time window selected based on ERF. It illustrates the spatial pattern of brain activation across four time-windows for each condition separately – faces and hands manipulation. C – Significant difference in Pseudo-Z between different conditions for the 330 - 450ms time window. OFA – occipital face area, pSTS – posterior superior temporal sulcus, FFA – Fusiform Face Area, PMC – primary motor cortex, IFG – inferior frontal gyrus, SMC – supplemental motor cortex.

Based on the observed components, we divided the visual trials in four time-windows and calculated Pseudo-Z maps for each condition for each window (Fig. 2B). During the first time-window (50-120ms) both faces and hand manipulations stimuli elicited activation in primary visual cortex in left and right hemisphere reflecting the first stage of visual information processing. Later, during 120-190ms the activation started to spread to early visual cortex involving ventral stream and dorsal stream areas in both hemispheres. In contrast, the activation for both conditions became more lateralized from 190 to 330ms with stronger involvement of right hemisphere where the activation continued extending to higher order visual areas. The most pronounced differences between two conditions (faces and hand manipulations) were detected during the last time window (330-450ms). Activation during face trials was located in the right lateral occipital (also known as occipital face area (Pitcher et al., 2011), posterior part of superior temporal cortex and inferior occipital and temporal (matching the location superior temporal visual area and fusiform face area), whereas hand manipulation trials elicited activation in postcentral and precentral cortical areas, and superior marginal gyrus, typically reported to be mirror neuron motor related areas (Grosbras et al., 2012).

PLS analysis confirmed the significance of observed differences in the spatial distribution of activation for the last time window (p < 0.001), while there were no significant differences during the previous time windows. The z-score distributions on the brain surface presented in Fig. 2, C, indicate the areas where faces and hands had different activation compared to each other. As expected, areas with higher activation during the fourth time window in face condition compared to hand manipulation included the ventral stream visual areas (occipital lateral area and fusiform area) and the superior temporal visual area. Whereas activation for hand manipulation condition was greater in supplementary and primary motor cortices and dorso-lateral frontal cortex.

#### 3.1.1. Synchronous oscillatory dynamics during faces and hand movements observation

The pseudo-Z contrast between faces and hand movement indicated peak pseudo-Z increase for faces in the ‘core system’ of face processing. Based on these results, we further investigated the temporal and spectral inter-trial oscillatory phase synchronization dynamics in both face and hand movements trials between these areas. In both conditions, synchronization was temporally more pronounced between 150 and 200ms in the alpha frequency, as illustrated in Fig. 3A and B columns. PLS analysis revealed statistically significant differences between the conditions (p < 0.05). The z-score distributions for all three connections in Fig 3C column indicate higher sPLV in the face condition which was the most pronounced in a laer period, between 200 and 300ms and in the alpha band. There are some strong z-score values in the baseline period, mostly at high frequency, suggesting that this connectivity estimation is noisy. Thus, interpretation should only be focused on cluster z-scores instead of local peaks.

**Fig.3.**
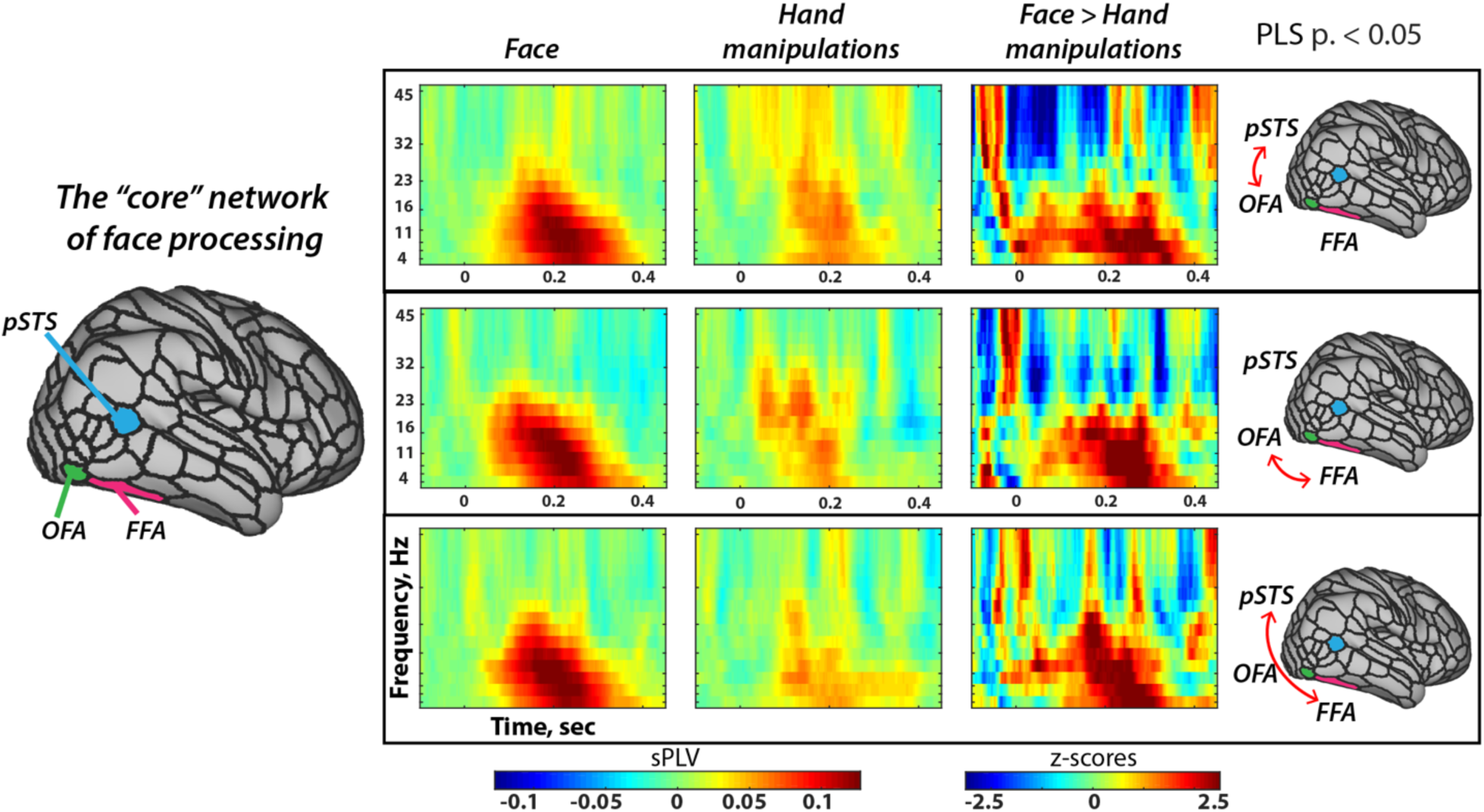
Dynamics of inter-trial phase synchronization between occipital face area (OFA), fusiform face area (FFA) and posterior superior temporal sulcus (pSTS) during faces and hand movements observation. A – sPLV dynamics during face condition between pSTS and OFA (first row), FFA and OFA (second row), pSTS and FFA (third row); B - sPLV dynamics during hand manipulation condition; C – PLS z-scores for each connection.

#### 3.1.2. Time frequency power changes in motor related areas

Pseudo-Z relative increase in motor related areas in the hand movement condition compared with the faces provided evidence of increased recruitment of the motor system. To characterize it in a spatio-temporal resolved manner, we analyzed the time and frequency activity of the areas classified as the motor system in the Glasser et al. atlas. The averaged TF representations across the motor areas in both conditions are illustrated in Fig. 4A and B. For both conditions, there is a low frequency spike after the onset of the stimuli. For the faces, however, the is mostly increase in power, especially in the beta, whereas, the hand movements, there is a pronounced decrease in power in the alpha and beta frequencies. These differences were statistically significant with PLS analysis (p-value <0.001). Based on the z-scores distribution in Fig.4C, power decrease in the beta frequency during the hand manipulations condition was contributing the most to the observed differences between the conditions.

**Fig.4.**
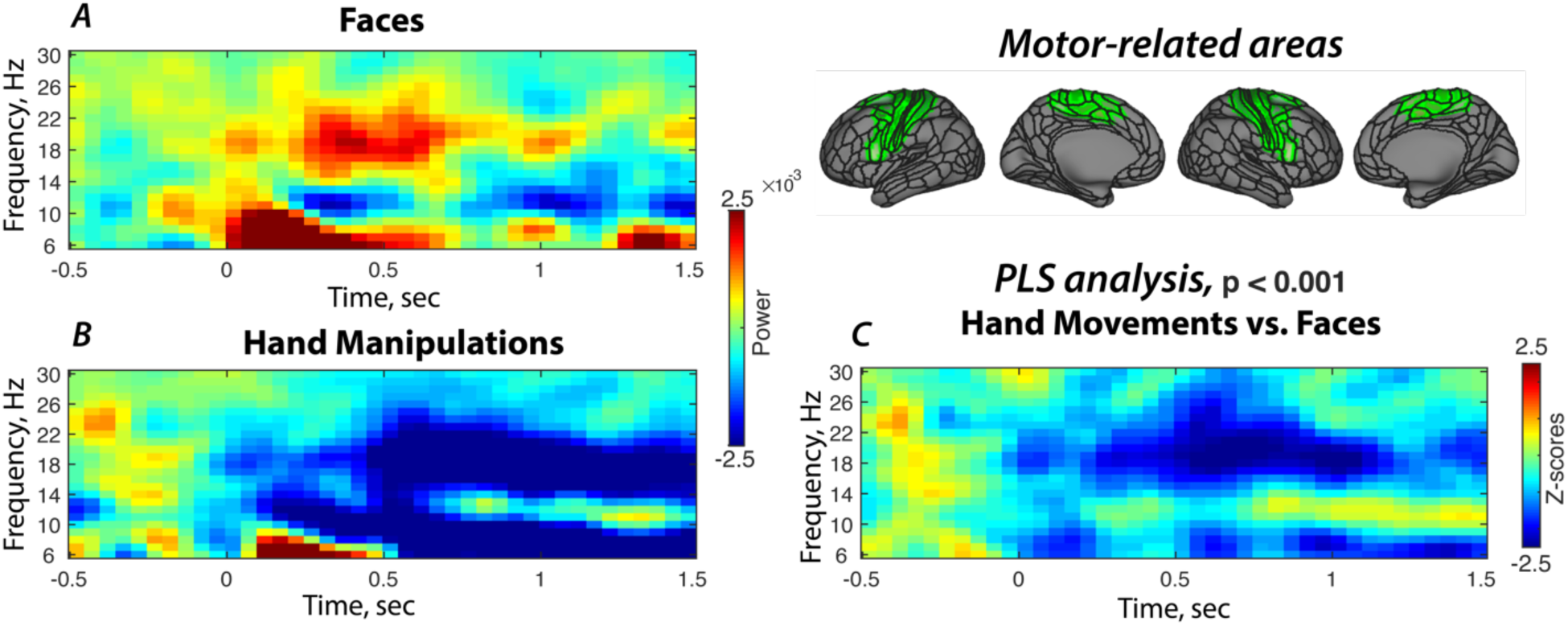
Spectral power dynamics in motor-related areas during faces and hand movements observation. A – time-frequency plot for face condition; B – time-frequency plot for hand manipulation condition; C – PLS z-scores illustrating the time and frequencies where the differences between conditions were the most pronounced.

### 3.2. Activation dynamics during listening to words and non-words

The analysis steps for mapping Pseudo-Z activation were further employed to investigate sensory processing of auditory stimuli by using trials with an onset of a word or a non-word sounds. Fig 5A shows an ERF averaged across all participants for left temporal MEG channels. Based on averaged ERFs, three time windows were chosen to perform source analyses for the auditory trials: 50-120ms (M100 component), 150-290ms (M200 component), 350-600ms (M400 component), which also are congruent with previous ERP studies of words and non-word stimuli (Cheng et al., 2014; Kaan, 2007).

**Fig.5.**
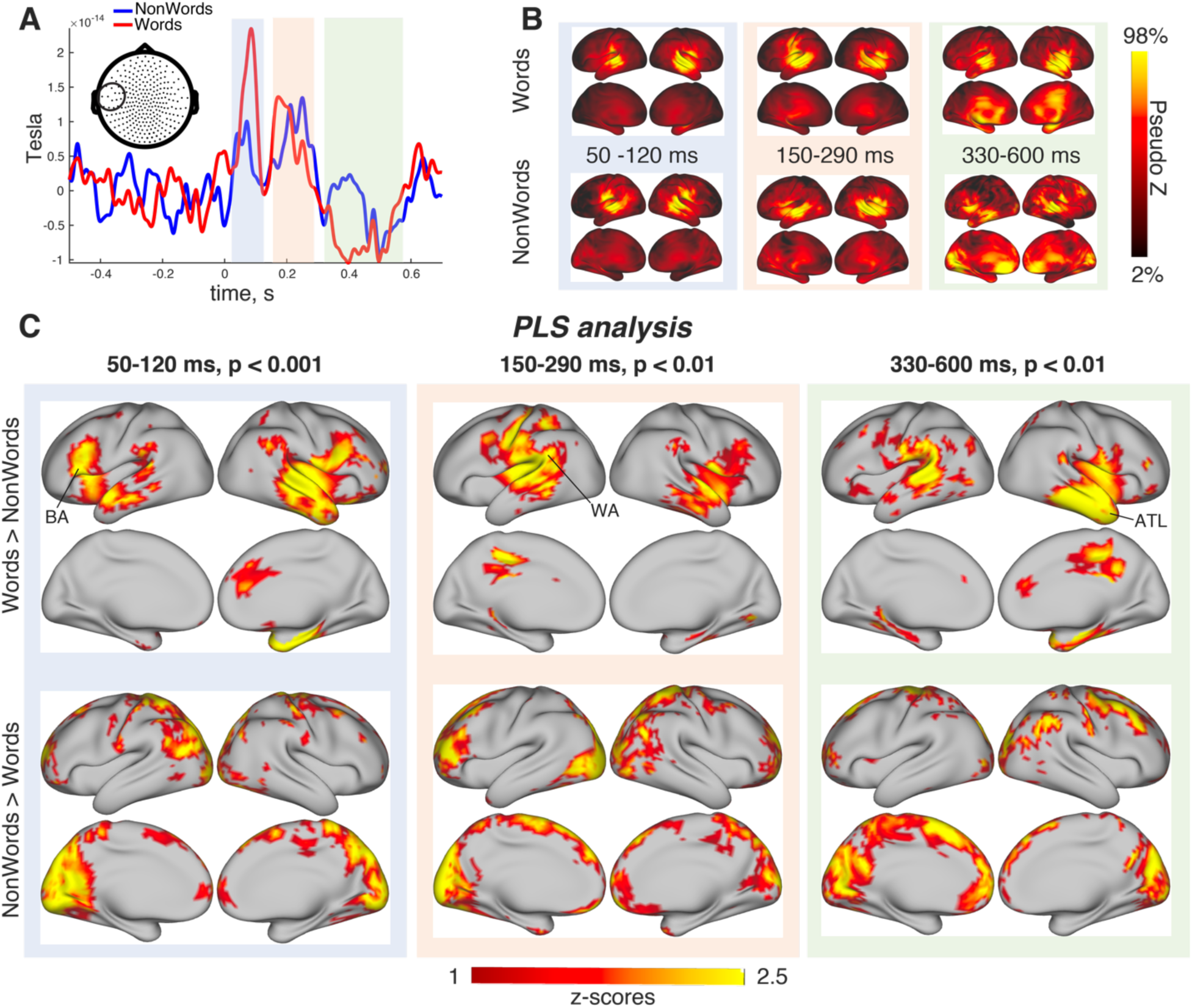
Differences in neuromagnetic dynamics between word and non-word auditory stimuli presented during the movie: A – averaged across left temporal sensors event related fields for both conditions. B – Pseudo-Z maps for each time window selected based on ERF. It illustrates the spatial pattern of brain activation across three-time windows for each condition separately – sounds composed of words and non-words. C – Significant difference in Pseudo-Z between different conditions for three time-windows: 50-120ms, 150-260ms and 330-600ms. BA – Broca’s area, WA – Wernicke’s area, ATL – anterior temporal lobe.

During the first time-window (50-120ms, Fig. 5B, blue panel) both word and non-words elicited activation in primary and secondary auditory cortices in both hemispheres. For the word condition, there was a noticeable higher activation in the right auditory cortex compared with non-word condition. This difference was further confirmed to be significant (p <0.001) using PLS activation (Fig. 3C, blue panel). Words stimuli elicited higher activation in the right anterior temporal pole (r-ATL), left inferior frontal gyrus (l-IFG), and the anterior part of the left superior temporal sulcus (l-STS), whereas non-words had higher activation in the left inferior parietal cortex and medial visual areas.

During the second time window (150-290ms, Fig. 3B, orange panel), word related activity in the STG spread along the superior temporal gyrus in the left hemisphere, and with less noticeable changes in the right hemisphere. In contrast, non-words stimuli activity expanded across the right superior temporal sulcus with fewer changes over time in the left hemisphere. PLS analysis on the pseudo-Z maps of the second time window revealed an overall significant contrast between the conditions (p < 0.01, Fig. 3C, orange panel). Higher activation was found in the l-STS including areas from temporal parietal junction (TPJ) for word trials, whereas for non-words higher activity was found in left lateral occipital cortex and left prefrontal cortex. Fig.3B, green panel illustrates the activation in left and right superior temporal gyrus for word stimuli during third time window (330-600ms). For non-word stimuli, the activation was more evident in right superior temporal area and left orbitofrontal areas. PLS analysis for the third time window revealed a significant contrast between conditions (p < 0.001, Fig. 3, C, green panel), with higher activation for word stimuli in left posterior part of the superior temporal cortex, TPJ and r-ATL. In non-words, higher activation compared to word stimuli was found in the calcarine sulcus, right middle and left superior frontal gyrus.

## 4. DISCUSSION

The present study aimed to demonstrate the feasibility of mapping previously characterized spatial patterns of brain activation and, in addition, their dynamics elicited by specific visual and auditory stimuli during a naturalistic movie watching paradigm in MEG. Although source space time-locked activation in neurophysiological studies are common, to our knowledge, this is the first time this type of analysis approach has been applied to a movie watching naturalistic paradigm, adapting a similar approach previously applied in fMRI (Lahnakoski et al., 2012). Using Pseudo-Z spatial distributions as an index of brain activation, we present an approach to map large-scale neurophysiological processing of visual (faces and hand movements) and auditory (words and non-words) stimuli over time windows defined by evoked potential fields (EPF). Such application in a naturalistic paradigm could be very beneficial when dealing with challenging populations. For example, it is well-known that in autism spectrum disorder face processing and mirror neurons system are disrupted, however, it remains challenging to study this population using neuroimaging, particularly individuals toward the low-functioning end of the spectrum. Watching a movie can make the brain recording enjoyable while valuable neuroimaging data can be obtained. Moreover, movie watching does not require the participant to follow strict task-related instructions, which is also a limiting factor in recording from clinical and child populations using common research protocols.

### 4.1. Face processing and hand movement observation during naturalistic viewing

Face processing involves a specialized distributed cortical network, predominantly, but not exclusively, lateralized in the right hemisphere (Barbeau et al., 2008; Pitcher et al., 2007). It starts at primary visual areas and progresses through a cortical hierarchy involving the occipital face area (OFA), the fusiform face area (FFA) and the posterior superior temporal sulcus (pSTS) face-selective area (Haxby and Gobbini, 2011; Pitcher et al., 2011; Turk-Browne et al., 2010). The last is reported to be sensitive to facial expressions and perception of body movements (Allison et al., 2000). Previous neurophysiological studies have reported M/N170, and M/N400f face-related components (Eimer, 2000; Harris and Aguirre, 2010; Itier et al., 2006; Liu et al., 2013), and in line with the literature, our results indicate that face related activity starts at primary visual areas, and over 450 milliseconds it expands to higher order and associative areas, mostly in the ventral stream. After Bonferroni correction, only in the latest temporal window 330-400ms face condition significantly differed from the hand-related movements condition. The areas with the highest peaks involved the right-OFA, right-FFA, right-pSTS, in the literature described as the ‘core system’ of face processing, and middle left-STS, described as part of the extended system and probably involved in comprehension face-related speech (Haxby et al., 2000). Based on these three peaks, we explored oscillatory phase synchrony between these areas. Our findings indicate significant contrast in inter-trial phase locking between faces and hand movements, with the most reliable differences located between 200 and 300ms in phase synchrony between the OFA, FFA and pSTS, although we did not find evidence of earlier synchronization between OFA and FFA, compared to pSTS.

While face perception recruits mostly ventral occipital-temporal areas, observation of hand movements in naturalistic viewing conditions activates mostly temporo-parietal areas (Avenanti et al., 2013). In particular, some of these areas have been associated with the mirror neurons system that activates when a person observes motor actions performed by others and is thought to facilitate the understanding of the ‘others’ actions (di Pellegrino et al., 1992; Mukamel et al., 2010).

The mirror neuron system is involved in a network of brain regions supporting theory of mind, which also includes the temporo-parietal junction (TPJ) (Brunet-Gouet and Decety, 2006; Vistoli et al., 2011), fronto-parietal regions including sensory-motor areas, and the dorsal and ventral frontal cortex (Avenanti and Urgesi, 2011; Romani et al., 2005). All these areas tend to be activated when observing actions performed by others (Arnstein et al., 2011; Grosbras et al., 2012; Mizuguchi et al., 2016) and with a right lateralization tendency (Bolognini et al., 2013; Stuss, 2001). Previously, the most pronounced effects of motor observation were reported to be at around 400ms (Balconi and Vitaloni, 2014; Casini et al., 2006; Reid and Striano, 2008). Similarly, in our study, significantly different activation in the hand movement condition compared to faces condition was found in the 330-450ms time window, possibly an overlapping period of late face processing and vicarious hand movement perception. The areas that differed the most compared to the face condition were located in somatosensory, primary and supplementary motor areas, dorsal prefrontal cortex, and inferior frontal cortex. Also, ventral prefrontal cortex increased activation was found, however, deep areas are more prone to localization errors and precaution should be taken when interpreting the results (Nunes et al., 2019).

Previous electrophysiological studies reported event-related desynchronization (ERD) during hand movements, as well as while observing others’ hand movements (Erbil and Ungan, 2007; Puzzo et al., 2011) suggesting that the motor activation induced by movement observation elicits a similar beta power desynchronization as when movement is performed (Hobson and Bishop, 2016; Meyer et al., 2011; Streltsova et al., 2010). Our findings in time-frequency analysis also demonstrated significant beta desynchronization across motor-related areas when participants observed hand movements compared to faces.

### 4.2. Contrasting brain activation dynamics for auditory words and non-words

In our study, we compared brain activation between the onset of words and non-word sounds. Based on previous literature, sound processing starts at primary auditory areas in the Heschl’s gyrus and moves forward to associative auditory areas where words activate the left lateralized cortical language circuit (Friederici, 2012; Price, 2009). It involves a dorsal stream that includes the ventral central sulcus (vCS), premotor cortex and IFG and the ventral stream, encompassing the posterior and anterior superior temporal gyrus (STG), and the anterior temporal lobe (ATL) (Friederici, 2012; Rauschecker and Scott, 2009). Our results found significantly different activation in the three time-windows of interest. In the first 50-120ms window, there was higher activation for words in the left anterior STG and IFG, but right vCS and STG. Previously, Liebenthal et al., 2013, also reported early (80-100ms) dorsal stream activation during phonemic perception. In the second window between 120-290ms, for words, there was a left-lateralization involving vCS and posterior STG where Wernicke’s area is located (Price, 2009), and in the last window between 330-600ms, while Wernicke area remained more active in the words condition, increased activation was more pronounced in the right hemisphere involving the ATL and middle temporal gyrus (MTG). The ATL, and the MTG, have been reported to act as a semantic hub from where semantic meaning is retrieved (Visser et al., 2012). Although it is more left-lateralized, both sides play an important role as demonstrated in lesion and TMS studies (Pobric et al., 2010; Ralph et al., 2016). Moreover, the right ATL might be more specialized in context processing and in sensory-motor representations such as voice and face recognition (Lindell, 2006; Olson et al., 2013). Higher activations during the non-words presentation were found scattered around the central sulcus and occipital cortex, interestingly, these activations are very similar to the activations evoked by listening to bird chirp sounds (Lindell, 2006). These findings could indicate a shift-balance in multisensory processing.

## Conclusions

In this study, we presented a series of time-locked analyses of dynamic brain activation patterns for faces, hand-movements, words and non-words conditions while participants were watching a movie. To our knowledge, this is the first demonstration of time-locked neuromagnetic responses with milliseconds time resolution during naturalistic viewing using MEG. The experimental control is lower during complex naturalistic stimuli, but by using pseudo-Z as a proxy of activation, the brain activity not related to the time-locked events was greatly suppressed, and meaningful activation related to movie events were localized. Significant differences in activation patterns between conditions allowed to tease apart the involvement of different brain areas in processing particular stimuli, and further source analysis proved the involvement of motor-related areas in movement observation and the involvement of face processing areas through oscillatory phase synchronization. Our results obtained in complex visual and auditory conditions are highly concordant with previous classical fMRI and neurophysiological studies. Similarities are described in terms of the brain regions activated for specific classes of stimuli, supporting and validating this approach for studying time-resolved sensory processing in a naturalistic paradigm.

## References

Allison, T., Puce, A., McCarthy, G., 2000. Social perception from visual cues: Role of the STS region. Trends Cogn. Sci. https://doi.org/10.1016/S1364-6613(00)01501-1

Arnstein, D., Cui, F., Keysers, C., Maurits, N.M., Gazzola, V., 2011. -Suppression during Action Observation and Execution Correlates with BOLD in Dorsal Premotor, Inferior Parietal, and SI Cortices. J. Neurosci. https://doi.org/10.1523/jneurosci.0963-11.2011

Avenanti, A., Candidi, M., Urgesi, C., 2013. Vicarious motor activation during action perception: beyond correlational evidence. Front. Hum. Neurosci. 7, 185. https://doi.org/10.3389/fnhum.2013.00185

Avenanti, A., Urgesi, C., 2011. Understanding “what” others do: Mirror mechanisms play a crucial role in action perception. Soc. Cogn. Affect. Neurosci. https://doi.org/10.1093/scan/nsr004

Balconi, M., Vitaloni, S., 2014. N400 effect when a semantic anomaly is detected in action representation. a source localization analysis. J. Clin. Neurophysiol. https://doi.org/10.1097/WNP.0000000000000017

Barbeau, E.J., Taylor, M.J., Regis, J., Marquis, P., Chauvel, P., Liégeois-Chauvel, C., 2008. Spatio temporal dynamics of face recognition. Cereb. Cortex. https://doi.org/10.1093/cercor/bhm140

Bolognini, N., Rossetti, A., Convento, S., Vallar, G., 2013. Understanding Others’ Feelings: The Role of the Right Primary Somatosensory Cortex in Encoding the Affective Valence of Others’ Touch. J. Neurosci. https://doi.org/10.1523/jneurosci.4498-12.2013

Brunet-Gouet, E., Decety, J., 2006. Social brain dysfunctions in schizophrenia: A review of neuroimaging studies. Psychiatry Res. - Neuroimaging. https://doi.org/10.1016/j.pscychresns.2006.05.001

Casini, L., Romaiguère, P., Ducorps, A., Schwartz, D., Anton, J.L., Roll, J.P., 2006. Cortical correlates of illusory hand movement perception in humans: A MEG study. Brain Res. https://doi.org/10.1016/j.brainres.2006.08.124

Centeno, M., Tierney, T.M., Perani, S., Shamshiri, E.A., StPier, K., Wilkinson, C., Konn, D., Banks, T., Vulliemoz, S., Lemieux, L., Pressler, R.M., Clark, C.A., Cross, J.H., Carmichael, D.W., 2016. Optimising EEG-fMRI for Localisation of Focal Epilepsy in Children. PLoS One 11, e0149048. https://doi.org/10.1371/journal.pone.0149048

Chang, W.-T., Jääskeläinen, I.P., Belliveau, J.W., Huang, S., Hung, A.-Y., Rossi, S., Ahveninen, J., 2015. Combined MEG and EEG show reliable patterns of electromagnetic brain activity during natural viewing. Neuroimage 114, 49–56. https://doi.org/10.1016/j.neuroimage.2015.03.066

Cheng, X., Schafer, G., Riddel, P.M., 2014. Immediate auditory repetition of Words and Nonwords: An ERP study of lexical and sublexical processing. PLoS One. https://doi.org/10.1371/journal.pone.0091988

Demirtaş, M., Ponce-Alvarez, A., Gilson, M., Hagmann, P., Mantini, D., Betti, V., Romani, G.L., Friston, K., Corbetta, M., Deco, G., 2019. Distinct modes of functional connectivity induced by movie-watching. Neuroimage 184, 335–348. https://doi.org/10.1016/J.NEUROIMAGE.2018.09.042

di Pellegrino, G., Fadiga, L., Fogassi, L., Gallese, V., Rizzolatti, G., 1992. Understanding motor events: a neurophysiological study. Exp. Brain Res. https://doi.org/10.1007/BF00230027

Eimer, M., 2000. Event-related brain potentials distinguish processing stages involved in face perception and recognition. Clin. Neurophysiol. https://doi.org/10.1016/S1388-2457(99)00285-0

Erbil, N., Ungan, P., 2007. Changes in the alpha and beta amplitudes of the central EEG during the onset, continuation, and offset of long-duration repetitive hand movements. Brain Res. https://doi.org/10.1016/j.brainres.2007.07.014

Friederici, A.D., 2012. The cortical language circuit: From auditory perception to sentence comprehension. Trends Cogn. Sci. https://doi.org/10.1016/j.tics.2012.04.001

Glasser, M.F., Coalson, T.S., Robinson, E.C., Hacker, C.D., Harwell, J., Yacoub, E., Ugurbil, K., Andersson, J., Beckmann, C.F., Jenkinson, M., Smith, S.M., Van Essen, D.C., 2016. A multi-modal parcellation of human cerebral cortex. Nature. https://doi.org/10.1038/nature18933

Glasser, M.F., Sotiropoulos, S.N., Wilson, J.A., Coalson, T.S., Fischl, B., Andersson, J.L., Xu, J., Jbabdi, S., Webster, M., Polimeni, J.R., Van Essen, D.C., Jenkinson, M., 2013. The minimal preprocessing pipelines for the Human Connectome Project. Neuroimage. https://doi.org/10.1016/j.neuroimage.2013.04.127

Grosbras, M.H., Beaton, S., Eickhoff, S.B., 2012. Brain regions involved in human movement perception: A quantitative voxel-based meta-analysis. Hum. Brain Mapp. https://doi.org/10.1002/hbm.21222

Harris, A., Aguirre, G., 2010. The effects of parts, wholes, and familiarity on face-selective responses in MEG. J. Vis. https://doi.org/10.1167/8.6.532

Hasson, U., Furman, O., Clark, D., Dudai, Y., Davachi, L., 2008. Enhanced Intersubject Correlations during Movie Viewing Correlate with Successful Episodic Encoding. Neuron 57, 452–462. https://doi.org/10.1016/J.NEURON.2007.12.009

Haxby, J. V., Hoffman, E.A., Gobbini, M.I., 2000. The distributed human neural system for face perception. Trends Cogn. Sci. https://doi.org/10.1016/S1364-6613(00)01482-0

Haxby, J. V, Gobbini, I.M., 2011. Distributed Neural Systems for Face Perception, in: Calder, A., Rhodes, G., Johnson, M., Haxby, J. (Eds.), Oxford Handbook of Face Perception. Oxford University Press, Oxford, pp. 93–110.

Hobson, H.M., Bishop, D.V.M., 2016. Mu suppression – A good measure of the human mirror neuron system? Cortex. https://doi.org/10.1016/j.cortex.2016.03.019

Honey, C.J., Thompson, C.R., Lerner, Y., Hasson, U., 2012. Not Lost in Translation: Neural Responses Shared Across Languages. https://doi.org/10.1523/JNEUROSCI.1800-12.2012

Itier, R.J., Latinus, M., Taylor, M.J., 2006. Face, eye and object early processing: What is the face specificity? Neuroimage. https://doi.org/10.1016/j.neuroimage.2005.07.041

Kaan, E., 2007. Event-Related Potentials and Language Processing: A Brief Overview. Lang.Linguist. Compass 1, 571–591. https://doi.org/10.1111/j.1749-818X.2007.00037.x

Kauppi, J.-P., Jääskeläinen, I.P., Sams, M., Tohka, J., 2010. Inter-subject correlation of brain hemodynamic responses during watching a movie: localization in space and frequency. Front. Neuroinform. 4, 5. https://doi.org/10.3389/fninf.2010.00005

Lahnakoski, J.M., Glerean, E., Salmi, J., Jääskeläinen, I.P., Sams, M., Hari, R., Nummenmaa, L., 2012. Naturalistic fMRI Mapping Reveals Superior Temporal Sulcus as the Hub for the Distributed Brain Network for Social Perception. Front. Hum. Neurosci. https://doi.org/10.3389/fnhum.2012.00233

Lankinen, K., Saari, J., Hari, R., Koskinen, M., 2014. Intersubject consistency of cortical MEG signals during movie viewing. Neuroimage 92, 217–224.https://doi.org/10.1016/j.neuroimage.2014.02.004

Lankinen, K., Saari, J., Hlushchuk, Y., Tikka, P., Parkkonen, L., Hari, R., Koskinen, M., 2018. Consistency and similarity of MEG- and fMRI-signal time courses during movie viewing. Neuroimage 173, 361–369. https://doi.org/10.1016/j.neuroimage.2018.02.045

Liebenthal, E., Sabri, M., Beardsley, S.A., Mangalathu-Arumana, J., Desai, A., 2013. Neural Dynamics of Phonological Processing in the Dorsal Auditory Stream. J. Neurosci. https://doi.org/10.1523/jneurosci.1511-13.2013

Lindell, A.K., 2006. In Your Right Mind: Right Hemisphere Contributions to Language Processing and Production. Neuropsychol. Rev. 16, 131–148. https://doi.org/10.1007/s11065-006-9011-9

Liu, J., Harris, A., Kanwisher, N., 2013. Stages of processing in face perception: An MEG study, in: Social Neuroscience: Key Readings. https://doi.org/10.4324/9780203496190

Lobaugh, N.J., West, R., McIntosh, A.R., 2001. Spatiotemporal analysis of experimental differences in event-related potential data with partial least squares. Psychophysiology 38, 517–30.

McIntosh, A.R., Lobaugh, N.J., 2004. Partial least squares analysis of neuroimaging data: Applications and advances. Neuroimage 23, 250–263. https://doi.org/10.1016/j.neuroimage.2004.07.020

Meeren, H.K.M., de Gelder, B., Ahlfors, S.P., Hämäläinen, M.S., Hadjikhani, N., 2013. Different Cortical Dynamics in Face and Body Perception: An MEG study. PLoS One. https://doi.org/10.1371/journal.pone.0071408

Meyer, M., Hunnius, S., Van Elk, M., Van Ede, F., Bekkering, H., 2011. Joint action modulates motor system involvement during action observation in 3-year-olds. Exp. Brain Res. https://doi.org/10.1007/s00221-011-2658-3

Mizuguchi, N., Nakata, H., Kanosue, K., 2016. The right temporoparietal junction encodes efforts of others during action observation. Sci. Rep. https://doi.org/10.1038/srep30274

Moiseev, A., Doesburg, S.M., Herdman, A.T., Ribary, U., Grunau, R.E., 2015. Altered Network Oscillations and Functional Connectivity Dynamics in Children Born Very Preterm. Brain Topogr. 28, 726–745. https://doi.org/10.1007/s10548-014-0416-0

Moiseev, A., Gaspar, J.M., Schneider, J.A., Herdman, A.T., 2011. Application of multi-source minimum variance beamformers for reconstruction of correlated neural activity. Neuroimage 58, 481–496. https://doi.org/10.1016/J.NEUROIMAGE.2011.05.081

Mukamel, R., Ekstrom, A.D., Kaplan, J., Iacoboni, M., Fried, I., 2010. Single-Neuron Responses in Humans during Execution and Observation of Actions. Curr. Biol. https://doi.org/10.1016/j.cub.2010.02.045

Nguyen, M., Vanderwal, T., Hasson, U., 2019. Shared understanding of narratives is correlated with shared neural responses. Neuroimage. https://doi.org/10.1016/j.neuroimage.2018.09.010

Nummenmaa, L., Glerean, E., Viinikainen, M., Jääskeläinen, I.P., Hari, R., Sams, M., 2012. Emotions promote social interaction by synchronizing brain activity across individuals. Proc. Natl. Acad. Sci. U. S. A. 109, 9599–604. https://doi.org/10.1073/pnas.1206095109

Nummenmaa, L., Saarimäki, H., Glerean, E., Gotsopoulos, A., Jääskeläinen, I.P., Hari, R., Sams, M., 2014. Emotional speech synchronizes brains across listeners and engages large-scale dynamic brain networks. Neuroimage 102, 498–509. https://doi.org/10.1016/J.NEUROIMAGE.2014.07.063

Nunes, A.S., Moiseev, A., Kozhemiako, N., Cheung, T., Ribary, U., Doesburg, S.M., 2019. Multiple constrained minimum variance beamformer (MCMV) performance in connectivity analyses. bioRxiv 567768. https://doi.org/10.1101/567768

Olson, I.R., McCoy, D., Klobusicky, E., Ross, L.A., 2013. Social cognition and the anterior temporal lobes: A review and theoretical framework. Soc. Cogn. Affect. Neurosci. https://doi.org/10.1093/scan/nss119

Oostenveld, R., Fries, P., Maris, E., Schoffelen, J.-M., 2011. FieldTrip: Open Source Software for Advanced Analysis of MEG, EEG, and Invasive Electrophysiological Data. Comput. Intell. Neurosci. 2011, 1–9. https://doi.org/10.1155/2011/156869

Pitcher, D., Walsh, V., Duchaine, B., 2011. The role of the occipital face area in the cortical face perception network. Exp. Brain Res. https://doi.org/10.1007/s00221-011-2579-1

Pitcher, D., Walsh, V., Yovel, G., Duchaine, B., 2007. TMS Evidence for the Involvement of the Right Occipital Face Area in Early Face Processing. Curr. Biol. https://doi.org/10.1016/j.cub.2007.07.063

Pobric, G., Jefferies, E., Lambon Ralph, M.A., 2010. Amodal semantic representations depend on both anterior temporal lobes: Evidence from repetitive transcranial magnetic stimulation. Neuropsychologia. https://doi.org/10.1016/j.neuropsychologia.2009.12.036

Price, C.J., 2009. The Year in Cognitive Neuroscience The anatomy of language: a review of 100 fMRI studies. Ann. N.Y. Acad. Sci. https://doi.org/10.1111/j.1749-6632.2010.05444.x

Puce, A., McNeely, M.E., Berrebi, M.E., Thompson, J.C., Hardee, J., Brefczynski-Lewis, J., 2013. Multiple faces elicit augmented neural activity. Front. Hum. Neurosci. 7, 282. https://doi.org/10.3389/fnhum.2013.00282

Puzzo, I., Cooper, N.R., Cantarella, S., Russo, R., 2011. Measuring the effects of manipulating stimulus presentation time on sensorimotor alpha and low beta reactivity during hand movement observation. Neuroimage. https://doi.org/10.1016/j.neuroimage.2011.05.071

Ralph, M.A.L., Jefferies, E., Patterson, K., Rogers, T.T., 2016. The neural and computational bases of semantic cognition. Nat. Rev. Neurosci. https://doi.org/10.1038/nrn.2016.150

Rauschecker, J.P., Scott, S.K., 2009. Maps and streams in the auditory cortex: Nonhuman primates illuminate human speech processing. Nat. Neurosci. https://doi.org/10.1038/nn.2331

Reid, V.M., Striano, T., 2008. N400 involvement in the processing of action sequences. Neurosci. Lett. https://doi.org/10.1016/j.neulet.2007.12.066

Robinson, S.E., Vrba, J., 1999. Functional neuroimaging by synthetic aperture magnetometry (SAM). Recent Adv. Biomagn.

Romani, M., Cesari, P., Urgesi, C., Facchini, S., Aglioti, S.M., 2005. Motor facilitation of the human cortico-spinal system during observation of bio-mechanically impossible movements. Neuroimage. https://doi.org/10.1016/j.neuroimage.2005.02.027

Schmuckler, M.A., 2001. What is Ecological Validity? A Dimensional Analysis. Infancy. https://doi.org/10.1207/S15327078IN0204_02

Schoffelen, J.-M., Hultén, A., Lam, N., Marquand, A.F., Uddén, J., Hagoort, P., 2017. Frequency-specific directed interactions in the human brain network for language. Proc. Natl. Acad. Sci. https://doi.org/10.1073/pnas.1703155114

Seymour, R.A., Rippon, G., Gooding-Williams, G., Schoffelen, J.-M., Kessler, K., 2018. Dysregulated Oscillatory Connectivity in the Visual System in Autism Spectrum Disorder. bioRxiv 440586. https://doi.org/10.1101/440586

Simony, E., Honey, C.J., Chen, J., Lositsky, O., Yeshurun, Y., Wiesel, A., Hasson, U., 2016. Dynamic reconfiguration of the default mode network during narrative comprehension. Nat.Commun. https://doi.org/10.1038/ncomms12141

Streltsova, A., Berchio, C., Gallese, V., Umilta’, M.A., 2010. Time course and specificity of sensory-motor alpha modulation during the observation of hand motor acts and gestures: A high density EEG study. Exp. Brain Res. https://doi.org/10.1007/s00221-010-2371-7

Stuss, D.T., 2001. The frontal lobes are necessary for theory of mind’. Brain. https://doi.org/10.1093/brain/124.2.279

Taylor, J.R., Williams, N., Cusack, R., Auer, T., Shafto, M.A., Dixon, M., Tyler, L.K., Cam-CAN Henson, R.N., 2017. The Cambridge Centre for Ageing and Neuroscience (Cam-CAN) data repository: Structural and functional MRI, MEG, and cognitive data from a cross-sectional adult lifespan sample. Neuroimage. https://doi.org/10.1016/j.neuroimage.2015.09.018

Turk-Browne, N.B., Norman-Haignere, S. V., McCarthy, G., 2010. Face-Specific Resting Functional Connectivity between the Fusiform Gyrus and Posterior Superior Temporal Sulcus. Front. Hum. Neurosci. https://doi.org/10.3389/fnhum.2010.00176

Vanderwal, T., Eilbott, J., Castellanos, F.X., 2019. Movies in the magnet: Naturalistic paradigms in developmental functional neuroimaging. Dev. Cogn. Neurosci. 36, 100600. https://doi.org/10.1016/J.DCN.2018.10.004

Vanderwal, T., Kelly, C., Eilbott, J., Mayes, L.C., Castellanos, F.X., 2015. Inscapes: A movie paradigm to improve compliance in functional magnetic resonance imaging. Neuroimage 122, 222–232. https://doi.org/10.1016/J.NEUROIMAGE.2015.07.069

Visser, M., Jefferies, E., Embleton, K. V., Ralph, M.A.L., 2012. Both the middle temporal gyrus and the ventral anterior temporal area are crucial for multimodal semantic processing: Distortion-corrected fMRI evidence for a double gradient of information convergence in the temporal lobes. J. Cogn. Neurosci. https://doi.org/10.1162/jocn_a_00244

Vistoli, D., Brunet-Gouet, E., Baup-Bobin, E., Hardy-Bayle, M.C., Passerieux, C., 2011. Anatomical and temporal architecture of theory of mind: A MEG insight into the early stages. Neuroimage. https://doi.org/10.1016/j.neuroimage.2010.09.015

